# Rejuvenation capacity of genetic versus human-optimized chemical reprogramming in cellular aging models

**DOI:** 10.1101/2025.11.27.690966

**Authors:** Omer Can Ergul, Tamer Onder

## Abstract

Cellular reprogramming with transient OSKM expression can reverse aging phenotypes, but genetic factor delivery introduces heterogeneous expression, reprogramming-associated stress, and barriers for therapeutic use. Small-molecule chemical reprogramming is an alternative, yet its performance relative to genetic approaches in human cells is unresolved. We directly compared a human-optimized chemical reprogramming protocol with doxycycline-inducible OSKM in progerin-induced aged fibroblasts and primary fibroblasts from donors over 85. Both reprogramming methods reduced senescence, mitochondrial ROS, and age-associated gene expression, with chemical reprogramming matching or exceeding OSKM efficacy. The two approaches, however, followed distinct trajectories. OSKM generated heterogeneous populations, including subsets acquiring pluripotency markers while others retained fibroblast identity. Chemical reprogramming produced uniform CD13-low populations without pluripotency marker induction. OSKM induced acute senescence that required a chase period to resolve, whereas chemical reprogramming lowered senescence during active treatment. In old fibroblasts, chemical reprogramming reversed multiple aging hallmarks while preserving fibroblast identity and avoiding telomerase activation. These results show that human-optimized chemical reprogramming can rejuvenate aged human fibroblasts with comparable efficacy to OSKM while generating more homogeneous outcomes and lower cellular stress, supporting small-molecule approaches as promising avenues for therapeutic rejuvenation.

## Introduction

Aging is the progressive decline of molecular order that compromises cellular function. This decline manifests as recurrent molecular and cellular alterations across tissues, formalized as the hallmarks of aging^1^. These include genomic instability, mitochondrial dysfunction, cellular senescence, and epigenetic alterations. Among these interconnected processes, epigenetic changes have attracted particular attention due to their potential reversibility and their ability to influence multiple downstream aging phenotypes.

Epigenetic structure defines how a single genome can establish and maintain distinct cellular identities. During aging, DNA methylation patterns undergo stochastic drift, histone modifications are aberrantly redistributed, and higher-order chromatin compartments deteriorate^2,3^. Importantly, they can be reversed through cellular reprogramming, indicating that aging phenotypes may be at least partially reversible^4^. Transient expression of the Yamanaka transcription factors (OCT4, SOX2, KLF4, c-MYC, collectively OSKM) can restore youthful cellular characteristics without inducing complete dedifferentiation when applied briefly^5^. Such partial reprogramming reduces senescence markers, decreases DNA damage, and restores mitochondrial function in both cultured cells and intact tissues^6-8^. However, transgene-based expression of reprogramming factors introduces substantial limitations. Viral or episomal delivery generates cell-to-cell heterogeneity in factor levels and triggers cellular stress responses. It also presents a narrow window between insufficient reprogramming and excessive pluripotency induction. These constraints are particularly pronounced in vivo, where spatial and temporal control of transgene expression remains challenging.

Small-molecule approaches to cellular reprogramming offer an alternative by modulating endogenous signaling pathways and chromatin regulators. Chemically defined cocktails can induce pluripotency and lineage conversion without genetic manipulation^9-12^. Chemical reprogramming offers several advantages. Compounds distribute homogeneously across cell populations, they can be administered in a time- and dose-dependent manner, and their effects are generally reversible. These properties should facilitate the optimization of partial reprogramming protocols. Recent studies have begun exploring chemical approaches for partial reprogramming and cellular rejuvenation, demonstrating improvements in aging markers in various model systems^13-15^. However, these efforts have largely relied on murine-optimized chemical cocktails applied to human cells or employed incomplete subsets of reprogramming protocols. Furthermore, direct comparisons between chemical and genetic reprogramming approaches under matched experimental conditions have not been performed, leaving key questions unresolved. Whether chemical reprogramming can reverse aging phenotypes in human cells as effectively as transcription factor-based approaches remains unclear. The extent to which the two methods operate through similar or distinct mechanisms is also unknown.

Primary human fibroblasts are a relevant cellular system to study aging and rejuvenation, exhibiting well-characterized age-associated phenotypes including elevated senescence-associated β-galactosidase (SA-β-gal) activity, increased mitochondrial reactive oxygen species (mtROS), accumulation of DNA damage foci, and progressive loss of heterochromatin markers^16-19^. However, the variability and modest magnitude of aging phenotypes in chronologically aged primary fibroblasts can complicate systematic optimization of reprogramming protocols. To establish a robust and reproducible aging model for initial testing and mechanistic comparison of reprogramming approaches, we employed stable overexpression of progerin. Progerin is the truncated lamin A variant that accumulates in Hutchinson-Gilford progeria syndrome (HGPS)^20^. Progerin expression recapitulates multiple hallmarks of physiological cellular aging, including nuclear envelope disruption, DNA damage accumulation, and premature senescence, in a pronounced and consistent manner^21^. This accelerated aging model enabled systematic optimization of chemical reprogramming parameters and direct comparison with genetic OSKM-mediated reprogramming. Following optimization in this tractable system, we validated our findings in primary human fibroblasts derived from old patients.

Here we applied the recently optimized human chemical reprogramming protocol described by Wang et al. (2025)^12^ to both progerin-expressing and naturally aged human fibroblasts, and directly compared its rejuvenation capacity and cellular dynamics with OSKM-mediated genetic partial reprogramming.

## Results

### Establishment and Characterization of a Progerin-Induced Cellular Aging Model

To systematically evaluate chemical reprogramming approaches for cellular rejuvenation, we first established a tractable model system that recapitulates key hallmarks of cellular aging. We generated a stable cell line by introducing a pBabe-Progerin-GFP fusion construct into young primary human fibroblasts (Fig. S1A). Progerin, the truncated lamin A protein that accumulates in Hutchinson-Gilford progeria syndrome (HGPS), has been well-documented to recapitulate multiple features of physiological aging at the cellular level.^22,23^ Overexpression of progerin in our model system induced characteristic aberrant nuclear morphology, including nuclear blebbing and irregular nuclear envelopes (Fig. S1B), consistent with previously reported HGPS phenotypes.

To comprehensively validate this aging model, we assessed multiple established hallmarks of cellular senescence and aging. Progerin overexpression significantly upregulated the expression of stress- and aging-associated genes (Fig. S1C), and dramatically increased the percentage of senescence-associated β-galactosidase (SA-β-gal)-positive cells (Fig. S1D) compared to WT lamin A expression. Additionally, progerin-expressing cells exhibited elevated mitochondrial reactive oxygen species (mtROS) levels (Fig. S1E). Consistent with age-related epigenetic alterations, we observed a significant reduction in heterochromatin marks, as evidenced by decreased H3K27me3 staining intensity (Fig. S1F). Together, these results confirm that progerin overexpression successfully recapitulates key molecular and cellular phenotypes of aging, providing a robust platform for evaluating rejuvenation strategies.

### Chemical Reprogramming Induces Rejuvenation in Progerin-Aged Fibroblasts Comparable to Genetic Reprogramming

Having established our aging model, we next evaluated the rejuvenation capacity of human-optimized chemical reprogramming. We employed the most recent human chemical reprogramming protocol described by Wang et al. (2025), which represents the first optimized cocktail specifically developed for human cells rather than adapted from murine protocols. This protocol comprises three sequential stages; we focused our studies on Stage 1, which consists of fourteen small molecules targeting key signaling and epigenetic pathways: 616452 (TGF-β inhibitor), CHIR99021 (GSK3β inhibitor), SAG (Smo receptor agonist), AM095 free acid (LPA1 receptor agonist), EPZ5676 (DOT1L inhibitor), WM-8014 (MOZ inhibitor), TTNPB (pan-RAR agonist), Ruxolitinib (JAK1/2 inhibitor), JNK-IN-8 (JNK inhibitor), DZNep (EZH2 inhibitor), VTP50469 (Menin-MLL inhibitor), A-485 (p300/CBP inhibitor), EZM0414 (SETD2 inhibitor) and AKT kinase inhibitor. We treated progerin-expressing aged fibroblasts with the Stage 1 cocktail for 2, 4, or 8 days, or transduced cells with a TET-inducible OSKM lentivirus and induced with doxycycline for the same durations, followed by a 4-day chase period in normal growth medium before analysis (Fig. 1A).

**Fig. 1.**
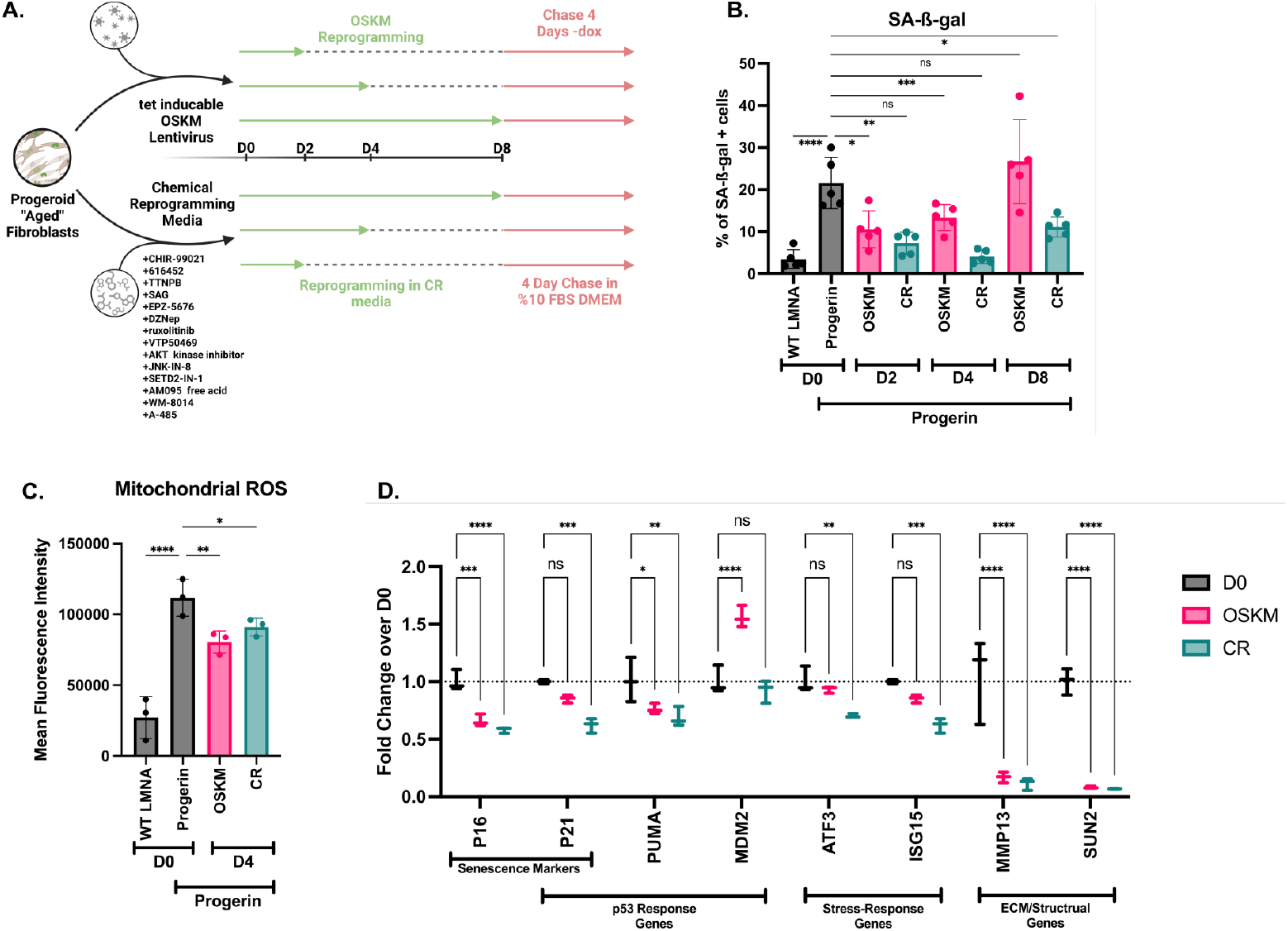
Chemical reprogramming induces rejuvenation in progerin-aged fibroblasts comparable to genetic OSKM reprogramming. (A) Experimental scheme showing progerin-expressing fibroblasts treated with Stage 1 chemical reprogramming cocktail or dox TET-inducible OSKM for 2, 4, or 8 days, followed by 4-day chase period. (B) Quantification of SA-β-gal-positive cells across treatment durations. (C) Mitochondrial ROS levels in progerin-expressing fibroblasts following 4 days of chemical or genetic reprogramming with 4-day chase, compared to Day 0 controls. (D) Expression of agingand stress-associated genes normalized to beta-actin in progerinexpressing fibroblasts following 4 days of chemical or genetic reprogramming with 4-day chase. Mean ± SEM, n=3 biological replicates. Two-way ANOVA with Šídák’s multiple comparisons test; *p<0.05, **p<0.01, ***p<0.001, ****p<0.0001.

To determine the optimal treatment duration, we first assessed senescence using SA-β-gal staining across all treatment conditions. While all chemical reprogramming durations significantly reduced the percentage of SA-β-gal-positive cells compared to untreated progerin-expressing controls (Day 0), we observed a non-linear dose-response relationship (Fig. 1B). The 2-day chemical treatment produced modest but significant senescence reduction, which was further enhanced with 4-day treatment. Notably, however, extension to 8 days of chemical exposure resulted in a partial reversal of this benefit, with SA-β-gal levels increasing relative to the 4-day treatment. This non-monotonic response suggests that prolonged chemical exposure may induce cellular stress, overwhelming the rejuvenation benefits. In contrast, 8-day genetic reprogramming using doxycycline-inducible OSKM resulted in elevated senescence markers relative to untreated Day 0 controls, likely reflecting prolonged reprogramming-induced stress. Based on these optimization experiments, we selected 4 days as the optimal chemical reprogramming duration for subsequent comparative analyses.

Using this optimized 4-day treatment protocol followed by a 4-day chase, we directly compared chemical reprogramming to genetic TET-inducible OSKM reprogramming. Both approaches achieved comparable reductions in mtROS levels relative to untreated progerin-expressing controls (Fig. 1C), indicating similar efficacy in restoring mitochondrial function. When we evaluated the expression of aging- and stressassociated genes, chemical reprogramming demonstrated slightly superior performance, achieving more pronounced downregulation of multiple senescence markers compared to OSKM-mediated genetic reprogramming (Fig. 1D). These results establish that human-optimized chemical reprogramming can effectively rejuvenate aged fibroblasts with efficacy matching or exceeding genetic approaches.

### Chemical Reprogramming Induces More Homogeneous Cellular Responses and Lower Reprogramming Stress Compared to Genetic OSKM

To gain deeper mechanistic insight into the distinct dynamics of chemical versus genetic reprogramming, we performed time-course analyses in young primary fibroblasts subjected to either OSKM or chemical reprogramming. Using flow cytometry with cell surface markers, we tracked the kinetics of cell state transitions at 2, 4, and 8 days of reprogramming (Fig. 2A).

**Fig. 2.**
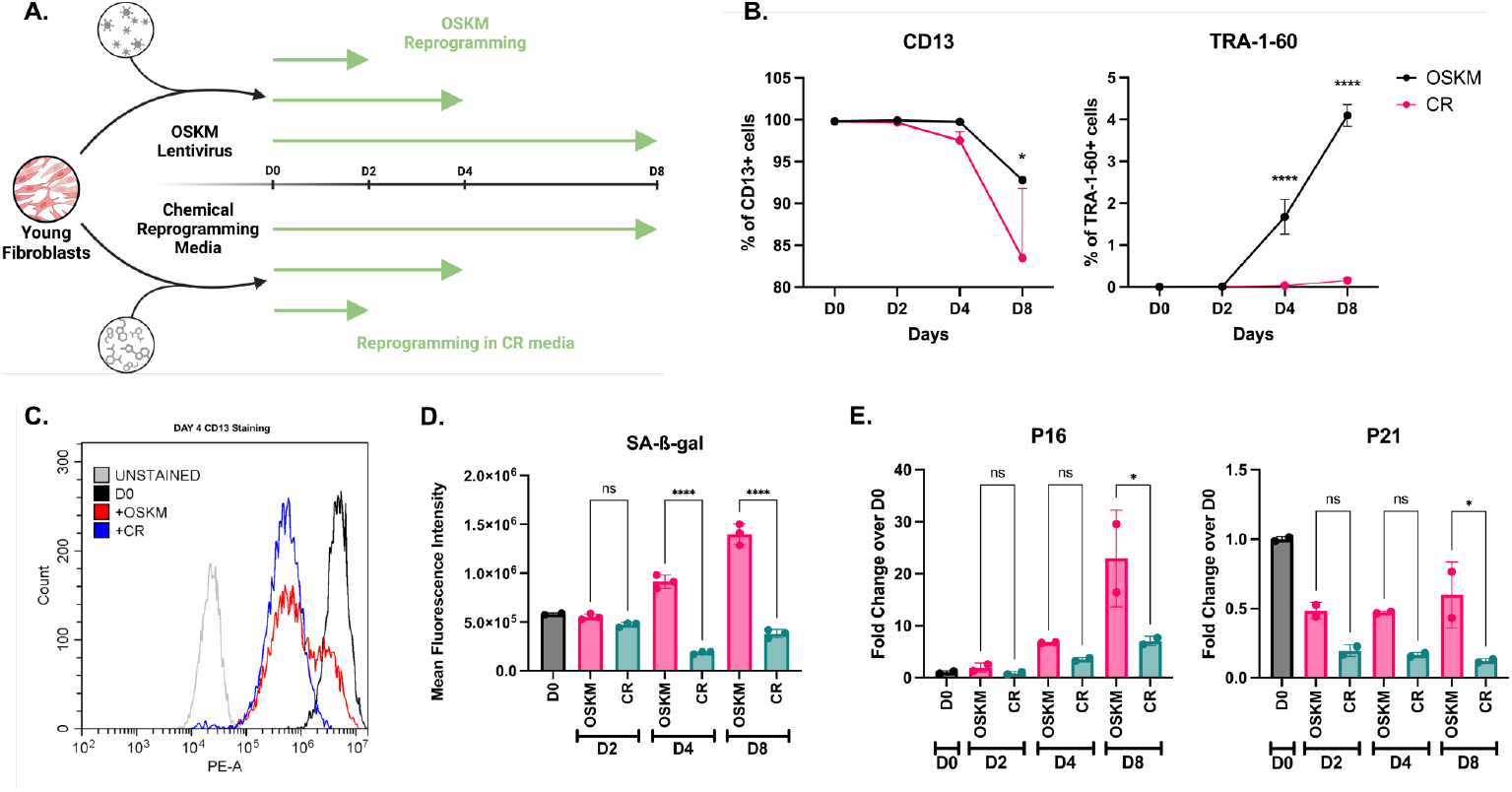
Chemical reprogramming induces homogeneous cellular responses with reduced stress compared to genetic OSKM reprogramming. (A) Experimental scheme showing young primary fibroblasts subjected to chemical or genetic OSKM reprogramming and analyzed at 2, 4, and 8 days. (B) Flow cytometry quantification of TRA-1-60+ (pluripotency marker) cells and CD13+ (fibroblast marker) showing greater and more uniform downregulation with chemical reprogramming. (C) Representative flow cytometry histograms of CD13 expression at Day 4 showing bimodal distribution in OSKM-treated cells versus uniform shift in chemically reprogrammed cells. (D) SA-β-gal quantification during reprogramming without chase period. (E) Expression of senescence markers p16 and p21 normalized to beta-actin following 4 days of reprogramming without chase. Mean ± SEM, n=3 biological replicates. Two-way ANOVA with Šídák’s multiple comparisons test; *p<0.05, **p<0.01, ***p<0.001, ****p<0.0001.

We assessed two key markers: TRA-1-60, a pluripotency-associated surface antigen, and CD13 (aminopeptidase N), a fibroblast identity marker.^24^ Remarkably, the two reprogramming approaches produced strikingly different outcomes. OSKM genetic reprogramming induced robust TRA-1-60 expression, with detectable percentages of TRA-1-60-positive cells detected at both day 4 and day 8 (Fig. 2B). In contrast, chemical reprogramming did not generate any detectable TRA-1-60-positive population, with levels remaining near baseline throughout the time course. Conversely, analysis of the fibroblast marker CD13 revealed chemical reprogramming induced a more dramatic loss of CD13 expression compared to OSKM reprogramming (Fig. 2C).

These seemingly contradictory observations, wherein OSKM cells acquire pluripotency markers while retaining fibroblast markers, while chemically reprogrammed cells lose fibroblast identity without acquiring pluripotency markers suggest fundamentally different reprogramming dynamics. OSKM genetic reprogramming appears to generate a highly heterogeneous cellular response, wherein the population separates into distinct subpopulations: some cells maintaining high CD13 expression characteristic of fibroblast identity, while others simultaneously acquire TRA-1-60 expression. This heterogeneity was further evident in flow cytometry histogram distributions at day 4 (Fig. 2D). While the chemically reprogrammed population shifted nearly uniformly toward CD13-low expression with minimal variance, the OSKM-treated population displayed a bimodal distribution, with approximately 35% of cells retaining CD13-high expression comparable to untreated fibroblasts (Day 0), and approximately 65% transitioning to CD13-low status. The chemical reprogramming approach, by contrast, induced a remarkably homogeneous response, with the vast majority of the population shifting coherently toward the CD13-low state without generating pluripotency marker-positive cells. These data indicate that chemical reprogramming drives cells through a more uniform, controlled trajectory that avoids the stochastic heterogeneity and potential progression toward pluripotency characteristic of OSKM-mediated reprogramming.

Given these distinct reprogramming dynamics, we next examined the stress responses associated with each approach. Reprogramming induction in primary cells has been extensively documented to trigger substantial cellular stress, including DNA damage responses and, paradoxically, senescence induction during the initial reprogramming phase.^25-27^ However, in the context of partial reprogramming, a reprogramming bout followed by a chase period, during which reprogramming factors are withdrawn has been shown to decrease cellular stress markers and reduce senescence.^5,6^ To test this, we quantified SA-β-gal activity immediately following reprogramming induction across the time course, without allowing a chase period (Fig. 2D). As expected, OSKM reprogramming without a chase period induced robust senescence, with SA-β-gal levels significantly elevated above baseline controls. Strikingly, chemical reprogramming showed no such requirement for a chase period, all time points of chemical treatment yielded senescence levels comparable to those achieved with a chase period. To determine whether the senescence induced by OSKM was attributable to the viral delivery method, we examined cells that received a mock viral transduction followed by 4 days of chemical reprogramming. While these virally transduced cells showed modestly elevated senescence relative to cells where chemical reprogramming was applied alone, the levels remained dramatically lower than OSKM-induced cells, suggesting that the difference observed between reprogramming modalities can’t be attributed to viral load alone (Fig. S2B). Gene expression analysis showed that both methods elevated p16 and reduced p21, but the magnitude differed substantially. Chemical reprogramming induced significantly less p16 while achieving a greater reduction in p21 (Fig. 2E).

Collectively, these findings demonstrate that chemical reprogramming achieves cellular rejuvenation through a fundamentally distinct mechanism characterized by homogeneous population dynamics and minimal activation of stress and senescence pathways, compared to the heterogeneous and stress-associated trajectory of OSKM-mediated genetic reprogramming.

### Chemical Reprogramming Effectively Rejuvenates Primary Aged Human Fibroblasts While Preserving Cell Identity

Having established the efficacy and mechanistic advantages of chemical reprogramming in controlled experimental systems, we sought to validate its translational potential in primary human cells derived from old donors. We isolated primary fibroblasts from tissue biopsies of four independent donors (>85 years old) and subjected these cells to partial chemical reprogramming using the optimized 4-day protocol (Fig. 3A).

**Fig. 3.**
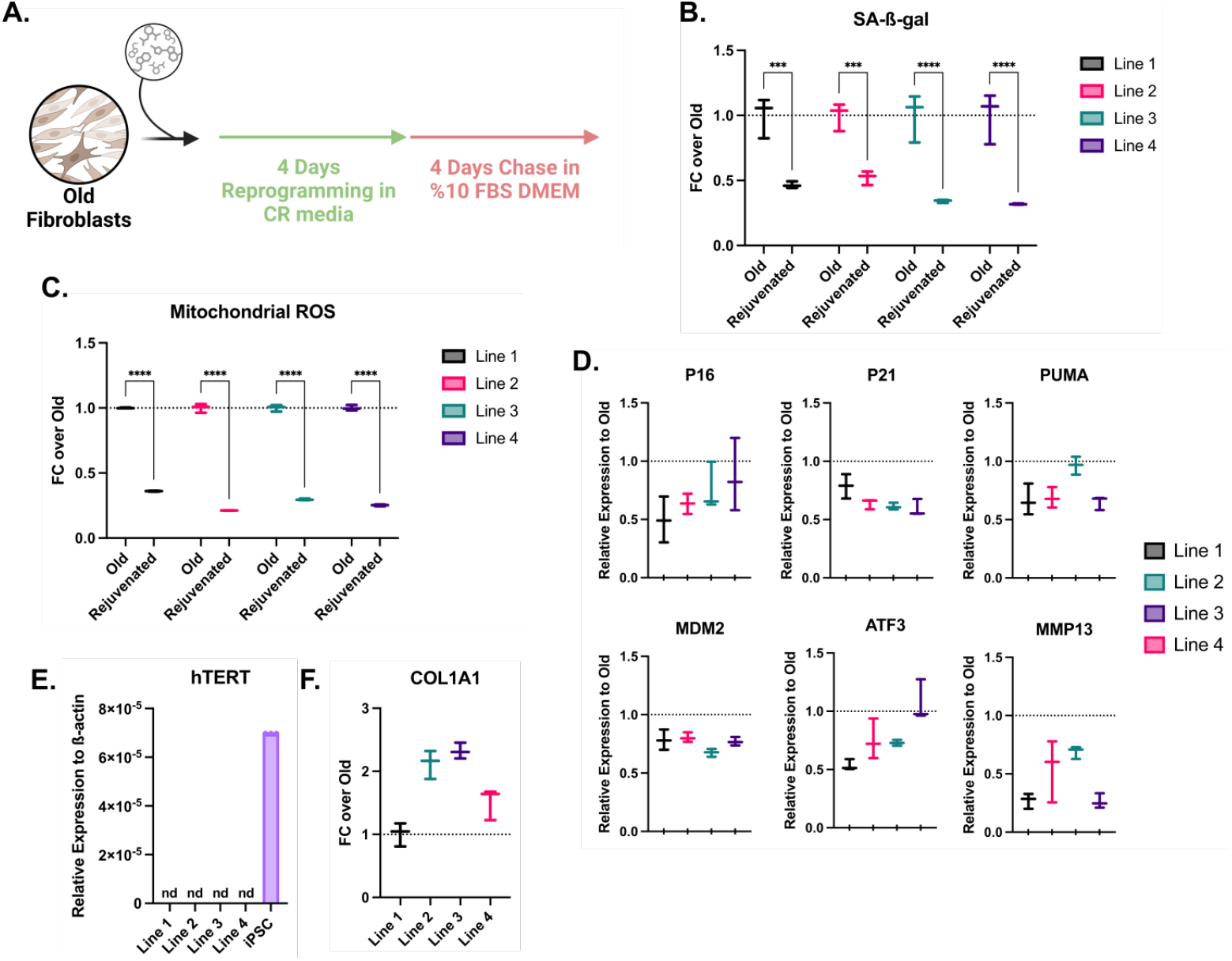
Chemical reprogramming effectively rejuvenates primary aged human fibroblasts while preserving cell identity. (A) Experimental scheme showing primary aged fibroblasts from four independent donors subjected to 4-day chemical reprogramming followed by 4-day chase. (B) SA-β-gal quantification showing reduced senescence across all donor lines. (C) Mitochondrial ROS levels following chemical reprogramming across four donor lines. (D) Expression of aging- and stress-associated genes normalized to beta-actin following chemical reprogramming across four donor lines. (E) TERT expression normalized to beta-actin following chemical reprogramming. (F) Expression of COL1A1 normalized to beta-actin following chemical reprogramming. Mean ± SEM, n=3 biological replicates per donor line across four independent donors. Two-way ANOVA with Šídák’s multiple comparisons test; *p<0.05, **p<0.01, ***p<0.001, ****p<0.0001.

First, we evaluated senescence levels following chemical reprogramming. The SA-β-gal-positive signal decreased significantly upon partial reprogramming across all donor lines (Fig. 3B), demonstrating consistent rejuvenation effects independent of donor-specific variability. Consistent with the decrease in senescence, we observed significant reductions in mtROS levels in all lines following chemical reprogramming (Fig. 3C), indicating that rejuvenation extended to metabolic and organellar function. We then evaluated transcriptional signatures of aging-associated genes following chemical reprogramming. Chemical treatment significantly downregulated multiple stress- and senescence-associated genes across all four primary fibroblast lines (Fig. 3D).

A critical concern in any rejuvenation strategy is the potential loss of cellular identity or uncontrolled proliferative capacity. We therefore assessed two key safety parameters: telomerase reverse transcriptase (TERT) expression, which could drive inappropriate cellular immortalization, and fibroblast-specific cell identity gene expression. Importantly, chemical reprogramming did not induce detectable TERT expression in any of the primary old fibroblast lines (Fig. 3E), indicating no activation of telomerase-mediated immortalization pathways. Furthermore, the expression of key fibroblast identity genes either remained stable or increased following chemical reprogramming (Fig. 3F), demonstrating that rejuvenation occurred without compromising cell-type-specific identity or function. In fact, an increase in genes related to cellular function has previously been shown to be in line with rejuvenation^8^.

Together, these results establish that human-optimized chemical reprogramming can effectively and safely rejuvenate naturally aged primary human fibroblasts, reducing multiple hallmarks of cellular aging while maintaining genomic stability and cell identity, essential prerequisites for therapeutic translation.

## Discussion

Here we directly compared human-optimized chemical reprogramming with genetic OSKM-mediated partial reprogramming to assess their capacity to reverse cellular aging hallmarks in human fibroblasts. Both approaches induced rejuvenation across multiple aging markers, including reduced senescence, decreased mtROS, and downregulation of age-associated genes. However, the two methods achieved these outcomes through markedly different cellular dynamics.

Previous studies have attempted chemical partial reprogramming in human fibroblasts, but these employed either murine-optimized chemical cocktails adapted to human cells or incomplete subsets of the full human reprogramming protocol^13-15^. None directly compared chemical and genetic reprogramming approaches under matched conditions. Our study is the first to apply the complete human-optimized chemical reprogramming cocktail and to systematically compare its rejuvenation efficacy and dynamics with OSKM-mediated genetic reprogramming. This direct comparison revealed fundamental differences in how the two modalities induce cellular reprogramming.

The most striking difference between methods was the homogeneity of cellular responses. OSKM reprogramming generated highly heterogeneous populations, with some cells downregulating the fibroblast marker CD13 while others retained high expression. Simultaneously, a subset acquired the pluripotency marker TRA-1-60. Chemical reprogramming, in contrast, induced remarkably uniform populationwide changes, with nearly all cells shifting coherently toward reduced CD13 expression without any detectable TRA-1-60-positive subpopulation. This difference is likely due to multiple factors. Viral or episomal delivery of reprogramming factors inherently generates cell-to-cell variability in transgene expression levels and stoichiometry. Beyond delivery, the reprogramming process itself is stochastic^28^. Individual cells encounter variable chromatin accessibility and signaling states that determine their response trajectories. Chemical compounds, by contrast, distribute uniformly and act on multiple targets simultaneously, potentially creating more coordinated cell-state transitions. These distinctions will likely be amplified in vivo, where uneven viral transduction, tissue structure, and local cellular environments would introduce additional variability.

The two approaches also differed substantially in cellular stress induction. OSKM expression triggered acute senescence responses, with elevated SA-β-gal activity within four days of factor induction. Chemical reprogramming reduced senescence markers even during active treatment and required no chase period. Gene expression analysis revealed that both methods induced p16 expression, though OSKM produced substantially higher levels. Despite elevated p16 expression, chemical reprogramming reduced SA-β-gal activity, revealing a disconnect between p16 transcription and the senescent phenotype. The elevated p16 in OSKM-treated cells is consistent with oncogene-induced senescence, a well-characterized barrier to reprogramming^30^. Some partial reprogramming studies have omitted c-MYC to reduce this effect^7^. Both approaches also reduced p21 expression, with chemical reprogramming achieving greater downregulation. The fact that chemical reprogramming achieved senescence reduction without a chase period, while OSKM required factor withdrawal, indicates distinct engagement with cellular stress pathways.

Our findings in naturally aged primary human fibroblasts confirm that chemical reprogramming can reverse aging hallmarks outside accelerated aging models. Across four independent donor lines, treatment reduced senescence, decreased mtROS, and downregulated age-associated genes without activating TERT or compromising fibroblast identity. The absence of telomerase activation and maintenance of lineage markers address key safety concerns for rejuvenation strategies.

Despite limitations of cell culture models, our findings establish that human-optimized chemical reprogramming can effectively reverse multiple aging hallmarks in human cells with efficacy comparable to genetic OSKM-mediated approaches. The use of the complete human reprogramming protocol on human cells, rather than adapted murine cocktails, represents a critical step toward translational relevance. The more homogeneous population dynamics and reduced cellular stress characteristic of chemical reprogramming address two major concerns for therapeutic translation: the risk of individual cells overshooting into pluripotency and the induction of stress-related pathways. These findings provide a foundation for developing small-molecule rejuvenation strategies as potential interventions for human aging and age-related diseases.

## Methods

### Cell Culture and Primary Fibroblast Isolation

Primary human dermal fibroblasts were isolated from skin biopsies obtained with written informed consent under Koç University Ethics Committee approval. Tissue samples were rinsed in PBS, minced into approximately 0.5-1 mm^3^ fragments, and placed in 6-well plates under sterile coverslips with DMEM supplemented with 15% fetal bovine serum (FBS) and 1% penicillin-streptomycin at 37°C with 5% CO2. Medium was changed at 24 hours and every 2-3 days thereafter. Fibroblast outgrowth typically appeared by days 4-6. Upon reaching approximately 80% confluence (days 10-14), cells were detached with 0.05% trypsin-EDTA, passaged, expanded, and cryopreserved for subsequent experiments. Young fibroblasts (derived from patients under 25 years old) were used for progerin model generation, while aged fibroblasts were isolated from donors over 85 years of age.

### Virus Production

HEK293T cells were seeded at 3.8 × 10^6^ cells per 10 cm dish in DMEM supplemented with 10% FBS and 1% penicillinstreptomycin. Cells were transfected with packaging plasmid (psPAX2 for lentivirus or pUMVC for retrovirus), envelope plasmid (VSV-G), and transfer plasmid using polyethylenimine (PEI) at a 1:3 DNA:PEI mass ratio. Following overnight incubation, medium was replaced with fresh DMEM + 10% FBS. Viral supernatants were collected at 48 and 72 hours, clarified by centrifugation at 3,000 × g for 10 minutes, and filtered through 0.45 µm membranes. For concentration, filtered supernatant was supplemented with polyethylene glycol 8000 (PEG-8000) to 10% (w/v), incubated at 4°C overnight, pelleted at 3,000 × g for 30 minutes, resuspended in ice-cold PBS, aliquoted, and stored at −80°C.

### Generation of Progerin-Expressing Cell Line

Young primary human fibroblasts were transduced with retroviral vectors encoding either wild-type lamin A (pBabe-LMNA-GFP-Puro, Addgene 17662) or progerin (pBabeProgerin-GFP-Puro, Addgene 17663) fused to GFP. Puromycin selection was applied until parallel non-infected control cells were eliminated. Selected cells were expanded in DMEM containing 10% FBS and cryopreserved for subsequent experiments.

### Genetic Partial Reprogramming

Fibroblasts were seeded at 3 × 10^4^ cells per well in 12-well plates. Cells were transduced with lentiviral vectors carrying doxycycline-inducible reprogramming factors as previously described^30^. The reprogramming cassettes were constructed by PCR-amplifying inserts from pSIN4-E2F-O2S (Addgene 21162) and pSIN4-CMV-K2M (Addgene 21164) and cloning into FuW-TetO-MCS (Addgene 84008), generating FUW-TETO-O2S and FUW-TETO-K2M constructs. Cells were co-transduced with these vectors along with FUW-M2-rtTA for doxycycline-inducible expression. Following transduction, cells were maintained under 5%O2 in DMEM with 10% FBS and 1% penicillinstreptomycin during the induction period. Doxycycline was added to induce reprogramming factor expression for the indicated durations (2, 4, or 8 days). Culture medium was replenished every two days. For partial reprogramming experiments with chase periods, following the indicated reprogramming duration, cells were cultured in DMEM with 10% tet-free FBS and 1% penicillin-streptomycin without doxycycline for 4 days before analysis.

### Chemical Partial Reprogramming

Chemical reprogramming was performed using a modified version of the Stage 1 protocol from the human-optimized chemical reprogramming system^12^. Cells were seeded at 3 × 10^4^ cells per well in 12-well plates in high-glucose DMEM with 15% FBS. After overnight attachment, cultures were switched to Stage 1 medium prepared in KnockOut DMEM containing 10% FBS, 10% knockout serum replacement (KSR), 1% GlutaMAX, 1% non-essential amino acids, 1% penicillinstreptomycin,LiCl (5 mM), nicotinamide (1 mM) and 50 µg/mL Vc2p. Small molecules were added as follows: CHIR-99021 (5 µM), 616452 (10 µM), TTNPB (2 µM), SAG (0.5 µM), EPZ-5676 (2 µM), DZNep (0.05 µM), ruxolitinib (1 µM), VTP50469 (0.5 µM), AKT inhibitor (1 µM), JNK-IN-8 (0.2 µM), SETD2-IN-1 (0.2 µM), AM095 (0.5 µM), WM-8014 (1 µM), and A-485 (0.5 µM). Cells were maintained under 5% O2. For rejuvenation experiments with chase periods, cultures were returned to standard growth medium (DMEM with 10% FBS) for 4 days to allow recovery before phenotypic assessment.

### Plate-based Senescence-Associated β-Galactosidase Staining

Cells were washed twice with PBS and fixed for 5 minutes at room temperature using 2% formaldehyde and 0.2% glutaraldehyde in PBS. After two PBS rinses, cells were incubated with freshly prepared X-gal staining solution (40 mM citric acid/sodium phosphate buffer pH 6.0, 5 mM potassium ferrocyanide, 5 mM potassium ferricyanide, 150 mM NaCl, 2 mM MgCl, 1 mg/mL X-gal in dimethylformamide) at 37°C for 14 hours in a non-CO2 incubator. Following staining, cells were washed twice with PBS and imaged. Cells were subsequently washed with methanol, air-dried, and stored protected from light. SA-β-gal-positive cells were identified by blue staining and quantified by bright-field microscopy.

### Flow cytometry-based Senescence-Associated β-Galactosidase Staining

Cells were treated with 100 nM bafilomycin A1 in %10 FBS DMEM for 1 hour at 37°C with 5% CO2 to elevate lysosomal pH. C12-FDG (5-dodecanoylaminofluorescein di-β-D-galactopyranoside) was then added directly to the medium at a final concentration of 15 µM and cells were incubated for an additional 2 hours at 37°C. Cells were washed twice with PBS, detached using 0.05% trypsin-EDTA, and neutralized with DMEM supplemented with 10% FBS. Following centrifugation at 1,000 × g for 5 minutes, cells were resuspended in ice-cold PBS and analyzed on a Beckman Coulter CytoFLEX flow cytometer. SA-β-gal activity was quantified as mean fluorescence intensity in the FITC channel.

### Mitochondrial ROS Measurement

Cells were washed twice with Hank’s Balanced Salt Solution (HBSS) without Ca2+/Mg2+, then incubated with 5 µM MitoSOX Red (TargetMol) in HBSS for 30 minutes at 37°C with 5% CO2. Following staining, cells were washed twice with PBS and detached with Accutase. Cells were centrifuged at 1,000 × g for 5 minutes and resuspended in PBS. Samples were analyzed on a Beckman Coulter CytoFLEX flow cytometer using the PE channel, and mitochondrial superoxide levels were quantified as median fluorescence intensity.

### RNA Isolation and Quantitative PCR

Total RNA was isolated using the NucleoSpin RNA kit according to manufacturer instructions. For cDNA synthesis, 500 ng to 1 µg RNA was combined with 0.5 µL of 10 mM dNTPs and 1 µL of 50 µM random hexamers, adjusted to 16.5 µL with nuclease-free water, heated at 65°C for 5 minutes, and chilled on ice. A reverse-transcription mixture containing 5 µL 5× First-Strand Buffer, 2 µL 0.1 M DTT, and 0.5 µL RNase inhibitor was added and incubated for 10 minutes at room temperature. Following addition of 1 µL MMLV reverse transcriptase, reactions were incubated at 37°C for 60 minutes and inactivated at 70°C for 15 minutes. cDNA was diluted to 100 µL with nuclease-free water.

Quantitative PCR was performed on a LightCycler 480 Instrument II using LightCycler 480 SYBR Green I Master in 20 µL reactions containing 10 µL master mix, 2 µL cDNA, 2 µL primer mix (each primer at 2.5 µM final concentration), and 6 µL nuclease-free water. Thermal cycling consisted of 40 cycles of 95°C for 30 seconds, 55°C for 30 seconds, and 72°C for 30 seconds. Relative expression was calculated using the 6.Ct method with normalization to housekeeping genes as indicated in figure legends.

### Flow Cytometry

Cells were detached with 0.05% trypsin-EDTA and neutralized with DMEM supplemented with 10% FBS. A total of 2 × 10^5^ cells were pelleted at 1,000 × g for 5 minutes and washed once with FACS buffer (4% FBS in PBS). Cells were resuspended in 100 µL staining buffer and incubated with 1 µL of fluorophore-conjugated antibody (1:100 dilution) for 30 minutes on ice in the dark. Primary antibodies used were PE-conjugated anti-human TRA-1-60-R (BioLegend 330610) and PE-conjugated anti-human CD13 (BioLegend 301704). Following incubation, cells were washed three times with staining buffer and resuspended in 300 µL FACS buffer. Samples with appropriate stained and unstained controls were analyzed on a Beckman Coulter CytoFLEX flow cytometer.

### Immunofluorescence Staining

Cells were seeded on 12 mm circular glass coverslips in 24-well plates. For imaging, cells were fixed in 4% paraformaldehyde for 30 minutes at room temperature, then washed three times with PBS for 5 minutes on an orbital shaker. For blocking and permeabilization, cells were incubated at room temperature for 60 minutes on an orbital shaker with 0.1% Triton X-100 and 5% goat serum in PBS. Primary antibodies were diluted in blocking buffer and incubated overnight at 4°C. Primary antibody used was rabbit anti-human H3K27me3 (Cell Signaling 9733). The following day, cells were washed three times with PBS for 5 minutes. Secondary antibodies (goat anti-rabbit Alexa Fluor 594, Invitrogen R37117) diluted in blocking buffer were applied for 1 hour at room temperature in the dark. Cells were washed three times in PBS for 5 minutes. Coverslips were mounted on microscope slides with VECTASHIELD Antifade Mounting Medium with DAPI and imaged on a Leica DMi8 confocal microscope. Fluorescence intensity was quantified using CellProfiler software.

### Statistical Analysis

All experiments were performed with three biological replicates unless otherwise stated. For experiments using primary aged fibroblasts (Figure 3), four independent donor lines were analyzed, with three biological replicates performed for each donor line. Data are presented as mean ± standard error of the mean (SEM). Statistical significance was determined by two-way ANOVA with Šídák’s multiple comparisons test using GraphPad Prism 10. Significance levels are indicated as follows: *p<0.05, **p<0.01, ***p<0.001, ****p<0.0001; ns, not significant.

## ACKNOWLEDGEMENTS

This work was funded by Turkish Institutes of Health (TUSEB) Grant no: 32810. The authors gratefully acknowledge the use of the services and facilities of the Koç University Research Center for Translational Medicine (KUTTAM).

## Supplementary Information

**Supplementary Figure 1.**
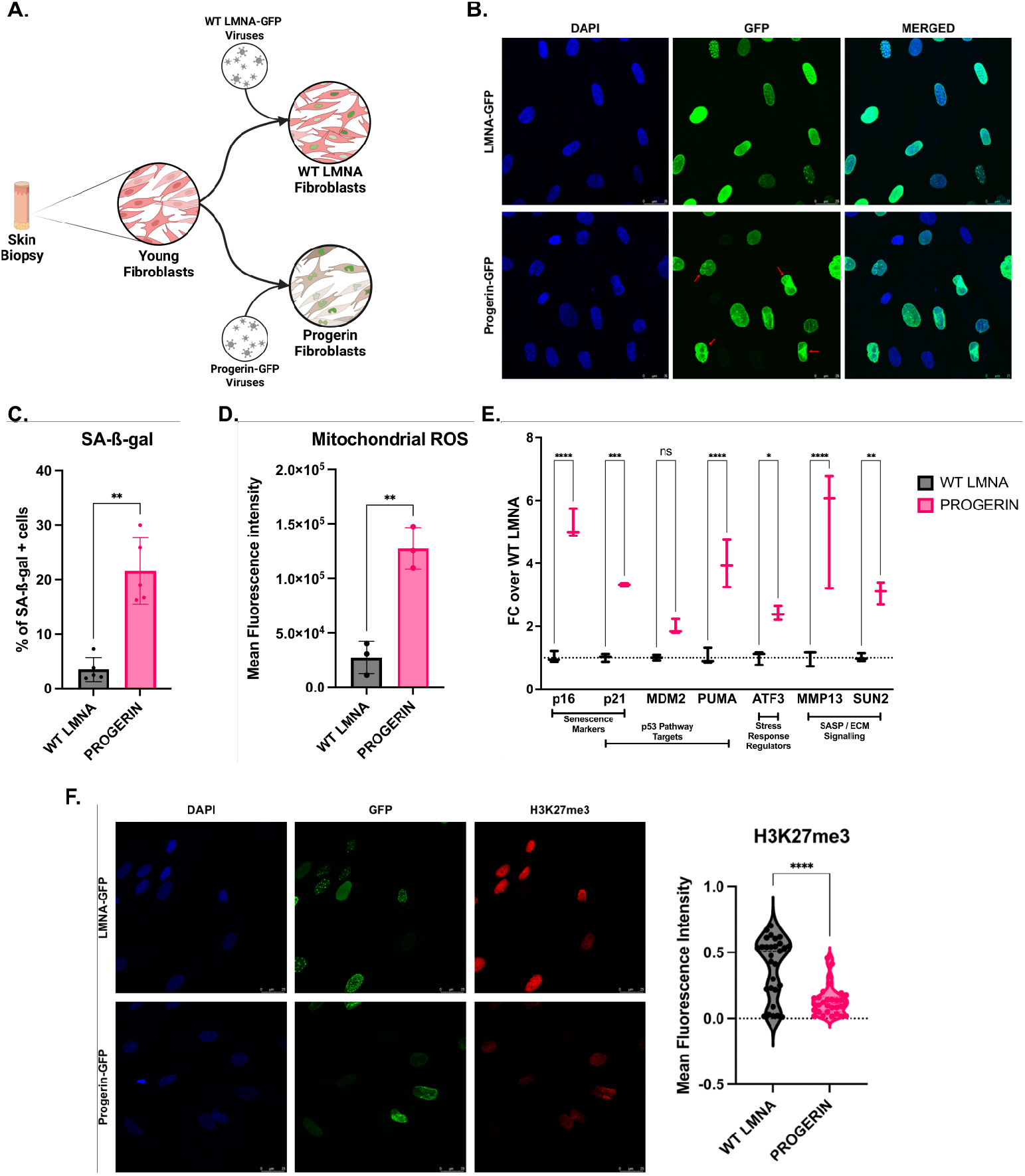
Progerin overexpression recapitulates cellular aging hallmarks in primary human fibroblasts. (A) Schematic of cell line generation of stable progerin-expressing and WT LMNA expressing cell lines. (B) Representative immunofluorescence images showing aberrant nuclear morphology in progerin-expressing cells (red arrows) compared to control fibroblasts. (C) SA-β-gal quantification showing increased senescence in progerin-expressing cells. (D) Mitochondrial ROS levels in progerin-expressing versus control fibroblasts. (E) Expression of aging- and stress-associated genes in progerin-expressing versus control fibroblasts. (F) H3K27me3 immunofluorescence quantification showing reduced heterochromatin marks in progerin-expressing cells. Mean ± SEM, n=3 biological replicates. Two-way ANOVA with Šídák’s multiple comparisons test; *p<0.05, **p<0.01, ***p<0.001, ****p<0.0001.

**Supplementary Figure 2.**
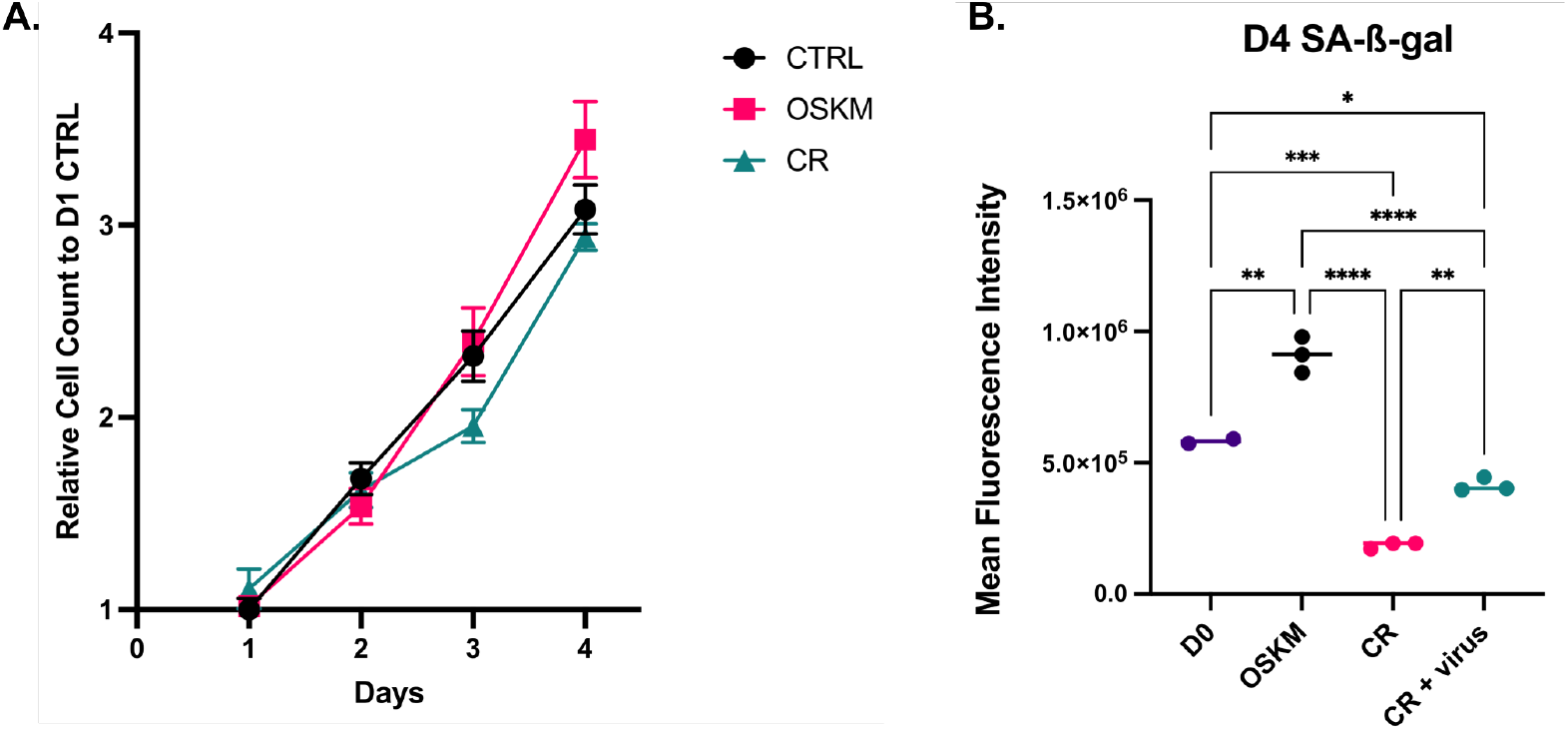
Cell viability and viral transduction effects during reprogramming. (A) CellTiter-Glo (CTG) luminescence assay showing relative cell viability measured daily during reprogramming. Young primary fibroblasts were left untreated (control), treated with Stage 1 chemical reprogramming cocktail (CR), or OSKM lentivirus. Viability was measured from Day 1 to Day 4. Data are normalized to Day 1 controls. (B) SA-β-gal quantification following 4 days of treatment in untreated controls (Day 0), OSKM-induced cells (OSKM), chemically reprogrammed cells (CR), and chemically reprogrammed cells with mock (rTTA) infection (CR + mock virus). Mean ± SEM, n=3 biological replicates. Two-way ANOVA with Šídák’s multiple comparisons test; *p<0.05, **p<0.01, ***p<0.001, ****p<0.0001.

